# Whole-Cortex Mapping of Common Genetic Influences on Depression and Social Deficits

**DOI:** 10.1101/500827

**Authors:** Alexander S. Hatoum, Andrew E. Reinberg, Harry R. Smolker, John K. Hewitt, Naomi P. Friedman

**Affiliations:** Institute for Behavioral Genetics, University of Colorado-Boulder; Department of Psychology and Neuroscience, University of Colorado-Boulder

**Author notes:** Correspondence concerning this article should be addressed to Alexander S. Hatoum, Institute for Behavioral Genetics, 447 UCB, University of Colorado, Boulder, CU 80309.

**Keywords:** Depression, Social, cortex, overlay, Twins, high-resolution

## Abstract

**Background:** Social processes are associated with depression, particularly understanding and responding to others, deficits in which can manifest as callousness/unemotionality (CU). Thus, CU may reflect some of the genetic risk to depression. Further, this vulnerability likely reflects the neurological substrates of depression, presenting biomarkers to capture genetic vulnerability of depression severity. However, heritability varies within brain regions, so a high-resolution genetic perspective is needed.

**Method:** In a sample of 258 same-sex twin pairs from the Colorado Longitudinal Twin Study (LTS), we developed a toolbox that maps genetic and environmental associations between brain and behavior at high resolution. We used this toolbox to estimate brain areas that are genetically associated with both depressive symptoms and CU. We then overlapped the two maps to generate coordinates that allow for tests of downstream effects of genes influencing our clusters.

**Results:** Genetic variance influencing cortical thickness in the right dorsal lateral prefrontal cortex (DLFPC) sulci and gyri, ventral posterior cingulate cortex (PCC), pre-somatic motor cortex (PreSMA), medial precuneus, left occipital-temporal junction (OTJ), parietal-temporal junction (PTJ), ventral somatosensory cortex (vSMA), and medial and lateral precuneus were genetically associated with both depression and CU. Split-half replication found support for both DLPFC clusters. Meta-analytic term search identified “theory of mind”, “inhibit”, and “pain” as likely functions. Gene and transcript mapping/enrichment analyses implicated calcium channels.

**Conclusions:** CU reflects genetic vulnerability to depression that likely involves executive and social functioning in a distributed process across the cortex. This approach works to unify neuroimaging, neuroinformatics, and genetics to discover pathways to psychiatric vulnerability.

---

As depression follows a normal distribution of risk across the population(1), relating depression to psychological features will better define pathways for addressing disorder vulnerability (2). Disruption in the ability to process social cues can lead to deficits in daily functioning and is often seen in depression(3). “Social Dimensions” is one of the main dimensions in the U.S. National Institute of Mental Health Research Domain Criterion (RDoC). Depressed individuals’ symptoms relate to specific facets of social behavior, namely, reasoning through others emotions(4–6), fitting under the subcategory of the “social dimensions” matrix, “understanding mental states.” Unsurprisingly, social deficits in theory of mind, the ability to understand others’ thoughts, are related to poor mentalizing/metacognition, or inability to understand the self. Further, theory of mind even predicts depression diagnosis above and beyond metacognition in behavioral studies(7).

An inability to understand and respond to others’ emotions may manifest as callousness/unemotionality (CU, 8). Although typically examined in the context of externalizing disorders (10, 11), CU has also been consistently associated with depression (10, 11). This association may arise because, while CU may reflect a disregard for others, it may also reflect an inability to empathize with others, perhaps due to poor theory of mind and metacognition about one’s own emotions. Consistent with this interpretation, CU has been related to mechanisms in social processing, like inability to understand others in development(8). Thus, poor social processing/CU may be a mechanism that sustains depression(3) or index the severity of depression(12). In either case, CU may reflect genetic influences on internalizing vulnerability(13), and brain mapping the overlap between depression and CU could help us determine whether cognitive or lower-order neurological systems are involved.

Multiple biological perspectives could advance our understanding of CU in depression. Family and genetic studies can estimate the relative importance of common genes and environments across two traits. Coinheritance between depression and CU is likely, as behavior genetics has established that depression is partially genetic in origin(14). Further, a recent genome-wide association analysis implicated over 150 genes in depression etiology(1), any of which could relate more specifically to social processing. However, while genetics is an excellent basis for studying psychiatric symptoms in the population, genes/variants and their downstream mechanisms are difficult to scrutinize(15).

In contrast to this lack of contextualization in genetic research, brain mapping integrates nicely onto other areas of biology (like transcriptomics(16)), thanks to the specificity gained when using high-resolution scanning coordinates. Here, we implement an integrative framework in which we directly map areas of the brain that represent the genetic overlap of CU and depression. Specifically, the goal of the current study is test whether CU captures some of the genetic vulnerability to depression; and localize the brain areas contributing to this vulnerability. These genetically associated brain areas can then be used with bioinformatic tools for mapping across different levels of the RDoC, such as RNA expression and biological pathway analyses, to expand our understanding of the coinheritance of CU and depression and find likely mechanisms of this behavioral vulnerability.

## Depression and CU in the Brain

Spatial brain mapping studies can localize where behavioral measures are associated with brain morphology. By overlaying neural correlates of depression with neural correlates from other measured behaviors, we gain specificity on areas associated with aspects of depression, like CU. While the neuroanatomical correlates of depression and CU have been studied extensively, this will be the first study examining the anatomical overlap between CU and depression.

The largest meta-analysis of neuroanatomical differences in depression to date used region of interest (ROI) measures of cortical thickness. It found that major depressive disorder (MDD) was associated with cortical thinning in the insula, anterior and posterior cingulate, and temporal gyri(17): areas key in salience(18), internal mentation(19), and switching between internal thought and executive control(20). However, this ROI approach does not consider how subcomponents of large areas may differentially relate to more specific facets of psychological phenomena; for example, anterior cingulate cortex shows differential gene expression and differential task activation across the ROI (21). One meta-analysis of voxel-based morphometry (VBM) found that MDD was associated with lower brain volume specifically in the rostral anterior cingulate cortex and the dorsolateral and dorsomedial frontal cortex(22).

For neuroanatomical correlates of CU, decreases in the volume of the rostral and dorsal cingulate cortex have been observed, overlapping spatially with the regions that have been identified for depression (23). Additionally, the rostral and dorsal anterior cingulate cortex areas that overlap between CU and depression were also found to distinguish suicidal cases from controls in another VBM study(24), giving some evidence that CU could represent a social severity dimension of depression.

## This Study

Using structural magnetic resonance imaging (MRI) data from 258 young adult twin pairs, we asked, where are the genes influencing the vulnerability to social deficits and depression influencing brain morphology? Do these morphological differences overlap? And, can we map a specific pattern and use this pattern to speculate further on mechanisms? To answer these questions, we used the methodology pictured in Figure 1 (a tutorial for this approach can be found on our github: https://github.com/AlexHatoum/Wild-Card-Toolbox). In step 1, we estimated the genetic and environmental association between depression and CU to evaluate the relative importance of shared inherited vulnerability. In step 2, we developed a toolbox that creates environmental and genetic brain maps for each trait. Rather than map standard beta coefficients (i.e., clusters associated with phenotypic variability), our procedure maps effect sizes for genetic and environmental variances (i.e., clusters associated with our traits via a genetic or environmental etiology), creating brain maps of genetic association between cortical thickness and the two behavioral traits. We estimate areas that represent the genetic vulnerability to CU and depression by overlaying the clusters from the separate depression and CU genetic maps onto one map. Finally, by integrating these brain maps with neuroinformatic tools in step 3, we can begin to characterize likely functions and specific molecular mechanisms of the genes influencing CU and depression, which is impossible in a standard biometrical design. Specifically, in step 3, we used MNI coordinates to align our genetically associated clusters with a meta-analytic database of effects across multiple fMRI and transcriptomic studies. Thus, our main analysis is the generation of genetically influenced brain map for depression and CU, and our follow-up analyses explore likely effects of this genetic variance implicated by this map by using high-resolution brain coordinates.

**Figure 1.**
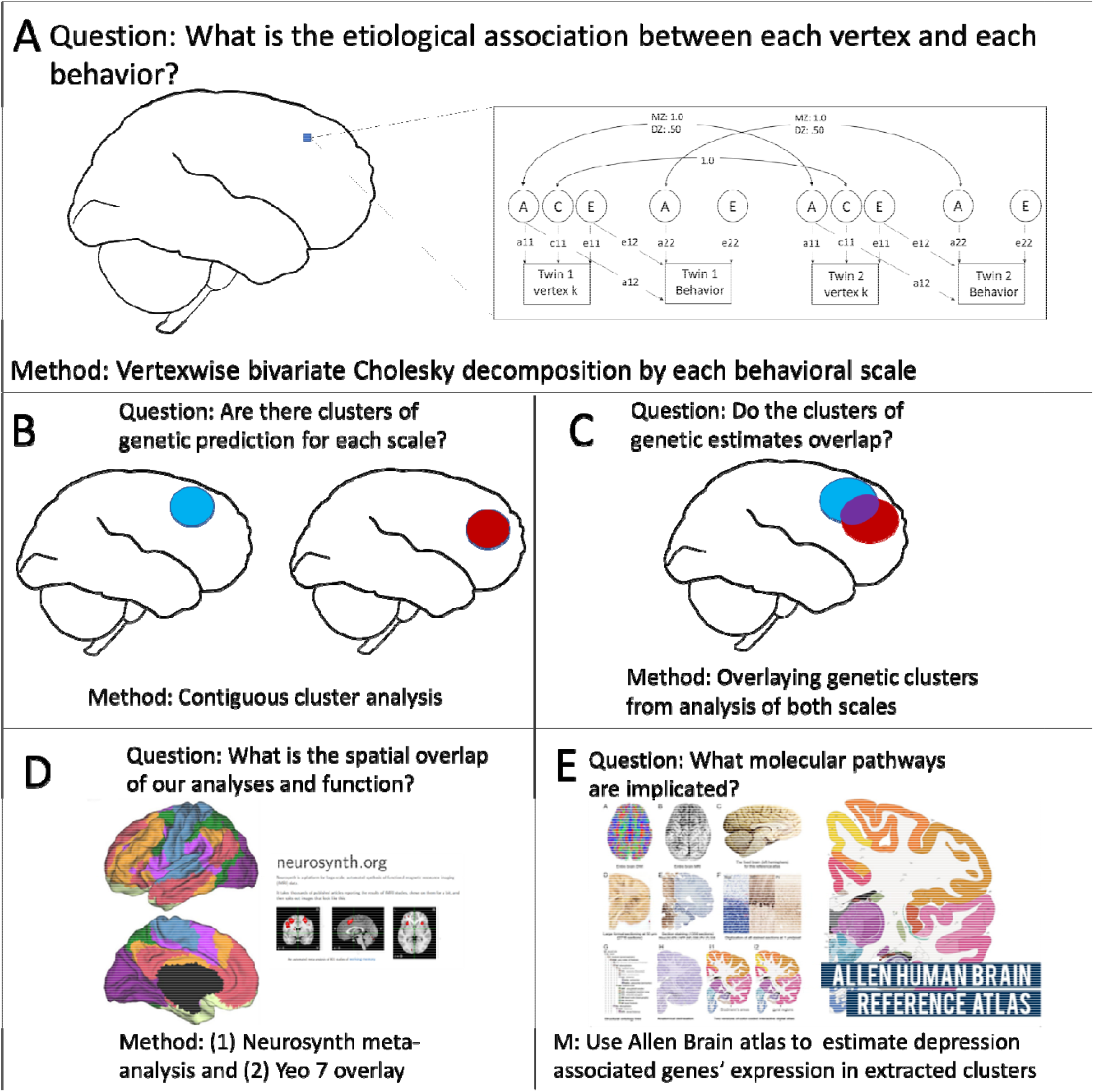
Five steps for whole cortex mapping by genetic association and follow up using informatic tools. Panel A: Additive genetic (A), Common environmental (C) and non-shared Environmental (E) Cholesky decomposition is used to find the etiological association of each vertex with each behavioral scale. Multiplication of standardized paths labeled 11 and 12 represents the phenotypic correlations predicted by additive genetic (Gr) and non-shared environmental (Er) influences, respectively. Panel B: Vertices whose associations with behavior are significant (p<. 05) and are part of a contiguous cluster of larger than 20 mm (cluster-extent correction) are estimated across the cortex surface separately by each trait and separately for A and E components. This procedure recovered 4 categories of clusters: additive genetic clusters influencing CESD, additive genetic clusters influencing ICU, non-shared environmental clusters influencing CESD, and non-shared environmental clusters influencing ICU. Panel C: Areas that represent significant conjunction of genetic association is created by overlaying the genetic clusters from CESD and ICU after cluster-extent correction. Panel D: The coordinates for overlap were transformed in MNI space and were used to map onto the Yeo 7 functional connectivity patterns and conduct meta-analytic term searches of likely associated functions. Panel E: Genes associated with depression in a large genome-wide association study were extracted from Neurosynth-gene/Allen Brain Atlas dataset to examine the expression of each of those genes in our clusters.

We conduct this analysis in a general population sample to include subsyndromal levels of depression and CU within a large enough sample to find patterns of coheritability between brain and phenotype. We chose high-resolution brain mapping because prior literature in neuroimaging genetics suggests vertex-wise approaches will more appropriately capture the individual differences patterns of genetic effects. In particular, common brain atlases used in anatomical research were derived agnostically to genes influencing individual differences and do not capture the specificity of the architecture of genetic effects on behavioral traits, as has been shown for language-related brain areas(25). Further, past work has shown there are differences in the genetic variance structure within and between commonly utilized ROIs; thus, measuring genetic variability in ROIs vs. vertices leads to relative differences in genetic variance effects between regions being overestimated(26) and more fine-grained metrics, such as voxel or vertex measures, are preferable to ROI approaches for making comparisons across the cortex for individual differences genetics(26). Notably for this study, it is these genetic individual differences patterns that are implicated in the mechanisms of psychopathology, requiring high-resolution coordinates to specify accurately. Finally, using high-resolution analysis and MNI coordinates allows for integration with functional and transcriptomics literature more broadly.

## Methods and Materials

### Sample

Participants were 258 same-sex twin pairs (135 monozygotic [MZ], and 123 dizygotic [DZ], 143 female pairs and 115 male pairs), aged 28 - 31 years (M = 28.7, SD = 0.6), recruited from the Colorado Longitudinal Twin Study (LTS). Twin pairs who had completed an ongoing neuroimaging study of neural substrates of executive functions and psychopathology and whose imaging data passed quality control were included. Two pairs were excluded due to cysts in the frontal cortex of one twin in each pair. More about the LTS can be found in the online methods.

### Structural MRI Scan

Images were acquired on a Siemens Trio 3 Tesla MRI scanner with 32-channel parallel imaging located at the University of Colorado Boulder. The total scanning session lasted 1 hour 25 minutes; the current analyses focus on gray matter structure, obtained with a high-resolution

T1-weighted Magnetization Prepared Gradient Echo sequence in 224 sagittal slices, with a repetition time (TR) = 2400 ms, echo time (TE) = 2.01 ms, flip angle = 8°, field of view (FoV) = 256 mm, and voxel size of 0.8 mm^3^.

### Behavioral Assessment

On the day of the scan, participants completed the Center for Epidemiological Studies-Depression (CESD) scale, a 20-item Likert scale assessing the frequency of past-week depression symptoms(27). We chose this measure because (1) tendencies toward an emotional vulnerability should manifest itself in higher frequency of depression, (2) we wanted to include subsyndromal levels of depression, and (3) this measure has shown reasonable stability across 10 years of longitudinal data(28).

Prior to the scanning session, participants completed an online questionnaire battery that included the Inventory of Callous and Unemotional traits (ICU)(11), a 24-item Likert questionnaire with three subscales: callousness (e.g., *The feelings of others are unimportant to me*), uncaring (e.g., *I do things to make others feel good*, reverse coded), and unemotional (e.g., *I do not show my emotions to others*). We used this scale as a measure of CU because it has been used to define clinical subtypes of conduct disorder in the past(29), the ICU total score relates to social and emotional processing(10), and, though the factor structure changes in adulthood, the scale retains a high internal consistency and predicts social, emotional, and depressive behaviors in individuals similar in age to our sample(29). We conducted all analyses with the ICU total scale, which is more reliable and normally distributed than the subscales, which were all highly intercorrelated (see Supplemental Table S1).

For the CESD and ICU, the dependent variable was the average item rating provided that at least 80% of the items were answered, multiplied by the number of items. To improve normality, both scales were then square-root transformed (see Supplemental Table 1).

### Data Analysis

All twins’ cortical thickness estimates were processed using a standard Freesurfer pipeline(30) (full description in online methods). Each vertex and psychopathology measure was residualized on brain mean thickness and sex prior to model estimation.

Behavioral genetic ACE models decompose phenotypic variance into three sources: Additive genetic (A; the sum of a large number of genetic variants), Common environmental (C; environmental influences that lead siblings to correlate), and non-shared Environmental (E; environmental influences that lead siblings to be uncorrelated). Because MZ twins share all their genes, their additive genetic influences correlated at 1.0; DZ twins share on average half their genes identical by descent, so their additive genetic influences correlate at 0.5. By definition, C effects correlate 1.0 and E effects correlate 0.0 for both types of twins.

To examine the genetic and environmental covariance between the psychopathology measures and brain measures, the standard ACE model for a single variable can be extended to multivariate analyses. To ensure that the estimated component covariance matrices are positive definite, they are expressed as the product of a lower triangular matrix and its transpose (Figure 1A). This is the Cholesky decomposition(31), which decomposes the phenotypic covariance between two measures into that explained by genes and environments. The genetic correlation (rG) of the two phenotypes equals (a11*a12)/√(a11^2^*(a12^2^ + a2^2^)).

#### Depression and CU coinheritance

To examine the etiological overlap between Depression and CU, we started by estimating their phenotypic overlap through a partial correlation analyses (accounting for sex and mean cortical thickness). We used a series of structural models to show that our association is specific to our measure of depressive symptom frequency and CU, rather than a broad psychiatric vulnerability (Supplemental Figure S1). Finally, we used a standard bivariate Cholesky decomposition to estimate the relative contribution of genes and environment to the overlap between the measures.

#### Discovery procedure for brain maps

A full diagram of the analysis plan is available in Figure 1. For each vertex, we estimated a separate Cholesky decomposition with the first variable being the vertex and the second being the CESD or ICU scale. We noticed substantial C variance across some areas of the cortex (Supplemental Figure S2) so we specified our Cholesky decompositions with a freed C path loading on the vertex but set the C cross path and specific C loading on the psychopathology variable to be zero, as there were no C effects on the CESD or ICU measures. We then computed the parameter representing the bivariate heritability, the phenotypic correlation predicted by the overlap in genetic influences (standardized a11*a12), at each vertex and projected it to a surface map in Freeview(32) to create whole-cortex heat maps of genetic effects on the brain-behavior association. From the generated whole-cortex map, we estimated clusters above significance for CESD and ICU, respectively, and then overlaid the CESD and ICU clusters.

To determine significant clusters for each disorder, we (1) estimated a chi-square difference test *p*-value for each Cholesky bivariate cross path, and (2) used vertex-wise cluster extent *p*-value correction of values below (.05) significance at a window of twice the original smoothing kernel (i.e. cluster extent threshold = 20 mm). We chose this procedure partially due to its practicality in integration with genetic estimates and to estimate clusters that were contiguous for follow-up analyses.

#### Split-half replication

To explore the replicability of our approach, we split our sample into halves by families (so that twin pairs would be kept together) by random draw (sample 1 *n* = 132 twin pairs, sample 2 *n* = 126 twin pairs) and ran the full analyses separately in each sample. In each half of the sample we used a conjunction minimum alpha of .05(33) and cluster extent correction of 20mm to define significant clusters. We then overlaid the clusters from the (1) the full sample analysis, (2) the analysis in sample 1, and (3) analysis in sample 2. Because the full sample was more conservative than either half, we wanted to use the criteria of significant overlap in all three analyses as our standard. i.e. a cluster must have been independently associated below the split half criteria in both half-samples and by a more conservative threshold with the full sample for us to have “high confidence” in its effect.

#### Transcripts, cell types, and functions associated with our genetic clusters

Using MNI coordinates, we examined the overlap of our clusters with other sources of data: (1) The Allen Brain atlas transcriptomic atlas and genome-wide association study (GWAS) results from the Psychiatric Genomics Consortium Depression mega-analyses(1), (2) Neurosynth meta-analytical database of functional activation across over 10,000 fMRI studies(34), and the (3) Yeo et al. 2011 7-network parcellation(35).

With the Allen Brain atlas, we took the list of associated genes from the psychiatric genomics consortium MDD GWAS gene-burden results(1) and used Allen brain atlas through Neurosynth gene(36) by downloading each gene image, renormalizing them across the cortex with FSL(37) and visualizing their expression. We excluded genes from the major histocompatibility complex, as these associations may be spurious due to long-range linkage disequilibrium (LD), and any genes not obtained through RNA arrays in the Allen Brain Atlas, leaving us with 100 genes. We put the expression values by each region in one matrix with k-means clustering. We used the elbow method (Supplemental Figure S3) see how many genetic clusters were recovered from our analyses. We then put the gene list of each cluster through the Reactome(38) pathway analysis database, using FDR to account for multiple comparisons.

With Neuro-synth, we entered our clusters from the discovery sample into Neuro-synth decoder to obtain “terms” that were most associated with functional activation across studies, as determined by a meta-analytic naïve Bayes classifier across over 10,000 fMRI studies(34). This analysis finds which of our coordinates most overlap with those found in the literature and which terms (fMRI patterns, tasks, or studied behaviors) are associated with those studies. We then identified which of these terms most commonly appeared across clusters (after filtering out nonspecific brain terms). Finally, we overlaid the coordinate of our clusters on the 7 resting-state networks from the Yeo parcellation(35) to identify to which networks the clusters belonged.

## Results

### Is CU Genetically Correlated with Depressive Symptoms?

We began by estimating the phenotypic, genetic, and environmental overlap between the depressive symptom frequency, measured by the CESD, and CU, measured by the ICU. Figure 2 shows the AE Cholesky decomposition. Based on the best fitting models for each univariate trait, C paths were not estimated (see Supplemental Table S2 for full model comparisons for each trait). We derived the genetic correlation between the two measures as *r*G= .40 (*p <.001*, see Supplemental Table S3 for genetic correlations between CESD and the ICU callous, uncaring, and unemotional subscales). The environmental association was not significantly greater than zero (*r*E=.04, *p*=.50). We concluded that the correlation between CU and depressive symptoms was due almost entirely to genetic covariance.

**Figure 2.**
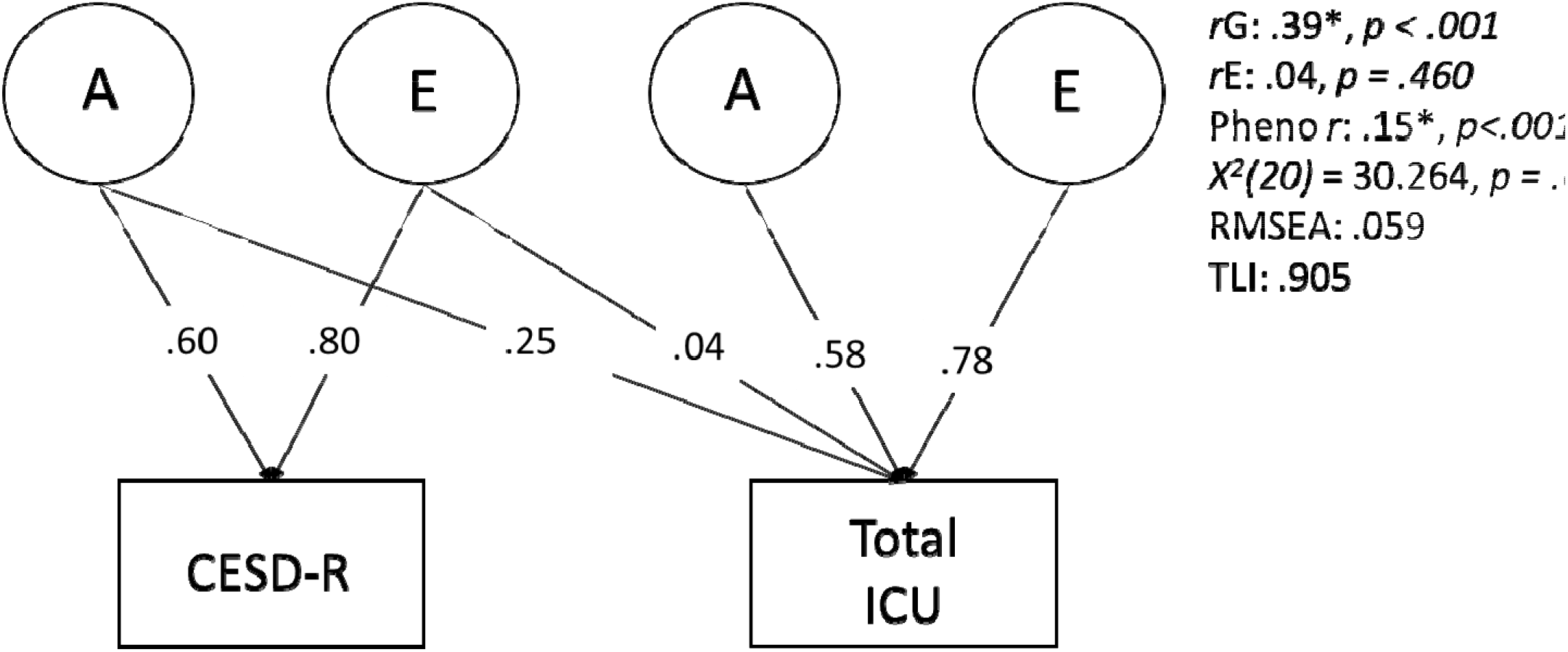
Additive genetic (A) and non-shared Environmental (E) Cholesky decomposition of the relationship between the Center for Epidemiological Studies Depression scale (CESD) and the Inventory of Callous and Unemotional traits (ICU). Numbers on arrows are standardized path estimates. Each task was residualized on sex and mean thickness prior to analysis. The derived genetic (*r*G), environmental (*r*E), and phenotypic (pheno *r*) associations are shown to the right of the path model. The model fit well by chi-square (*X^2^* (20) = 30.264, *p = .070* and RMSEA (.059). \**p*<.05, determined by 1-df chi-square difference test. Dotted lines indicate p>.05.

### Where are CU/Depressive Symptom Genetic Influences Related to Brain Morphology?

We created a map of areas where cortical thickness genetically correlated with CESD and ICU scores. We then overlaid the clusters from the two maps to discover regions that showed conjunction for genetic prediction.

As shown in Figure 3 and Table 1, we found that genetic influences on thicker cortex in the right dorsal lateral prefrontal cortex (DLFPC) sulci, the right pre-somatic motor cortex (PreSMA), left medial and lateral precuneus, occipital-temporal junction (OTJ), and temporoparietal junction (TPJ) were associated with both traits (i.e., these areas showed positive genetic associations above significance with both measures). We found genetic influences on thinner cortex in the right ventral posterior cingulate cortex (PCC), right medial precuneus, right DLPFC gyrus, and left ventral somatosensory cortex in the pathophysiology of both traits. Finally, split half-replication gave support for both right DLPFC areas in the same direction as discovered in the full sample (Supplemental Figure S4). Comparison to phenotypic maps (Supplemental Figure S6) showed that overlay regions discovered would have been qualitatively different without the genetic approach, as phenotypic areas did not overlap substantially with our genetic areas.

**Figure 3.**
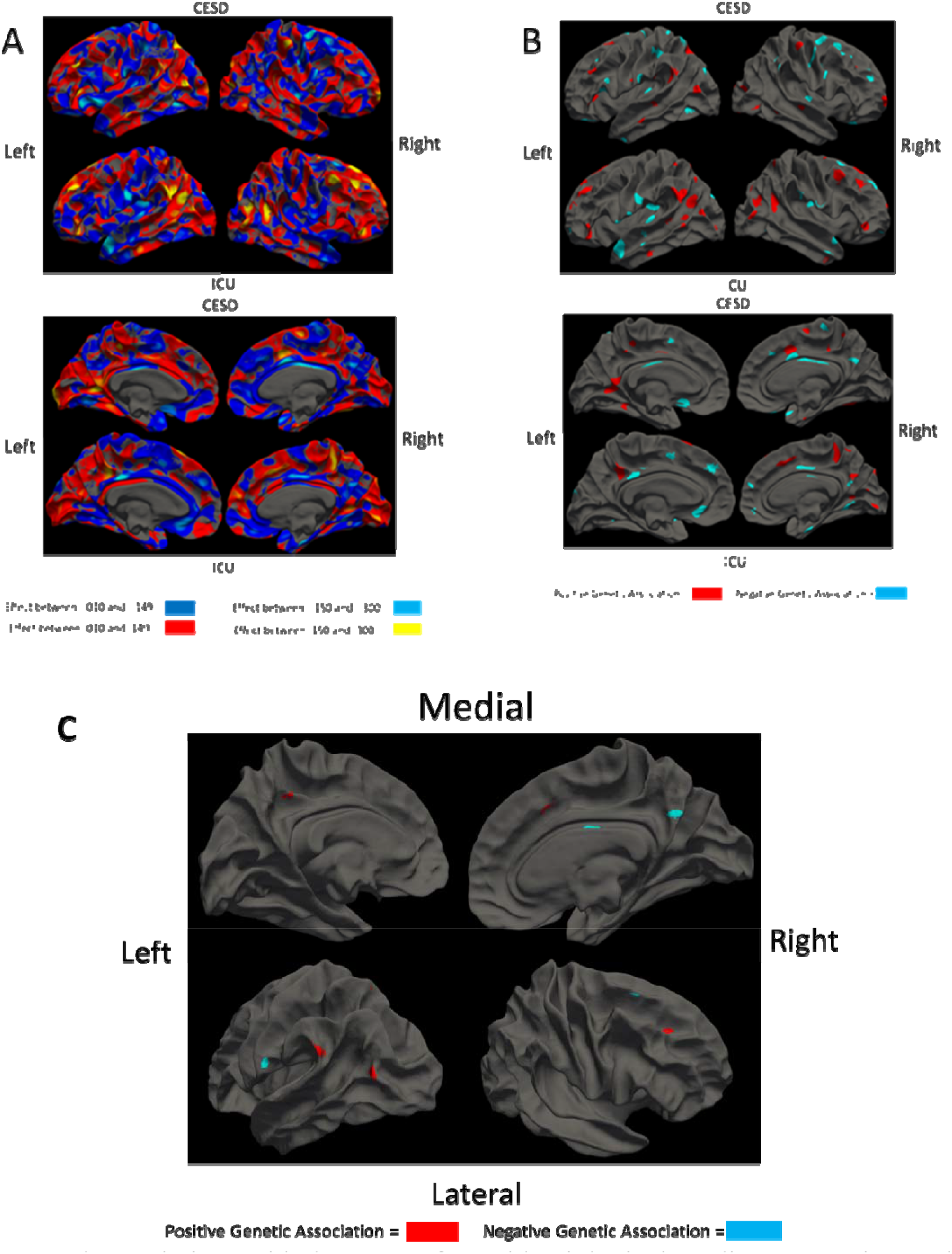
Neural associations with the Center for Epidemiological Studies Depression scale (CESD) and Inventory of Callous and Unemotional traits (ICU). Panel A depicts whole-cortex heat maps of the genetic association of each vertex with each behavioral measure as bivariate heritability. Panel B depicts *p*-values for genetic association between each vertex and each behavioral scale below correct significance (*p* < .05). Lateral views are on top and medial views below. These analyses correspond to those outlined by Figure 1 panel B. Panel C depicts overlap areas for our genetic clusters. These genetic clusters coordinates were used in all future analyses.

**Table 1.** Cluster Coordinates for Overlay Clusters in mm Space

| **Cluster** | **COG X** | **COG Y** | **COG Z** | **Number of Vertices** |
| --- | --- | --- | --- | --- |
| **L-Lateral Precuneus** | -14 | -67 | 57 | 6 |
| **L-Medial Precuneus** | -6.54 | -42.3 | 43.6 | 38 |
| **L-Occipital Junction** | -46.1 | -72.8 | 14.7 | 138 |
| **L-Temporal Junction** | -57.7 | -49.1 | 29.9 | 191 |
| **L-ventral SMA** | -60.8 | -16.7 | 23.9 | 73 |
| **R-DLPFC\*** | 23.5 | 32.6 | 35.2 | 61 |
| **R-Lateral Frontal\*** | 23 | 16.4 | 57.2 | 21 |
| **R-PCC** | 4.87 | -13 | 30.6 | 42 |
| **R-Posterior Precuneus** | 5.48 | -59 | 31.1 | 99 |
| **R-PreSMA** | 11.1 | 12.6 | 43.6 | 28 |

*Note.* Cluster coordinates for each of the overlay clusters discovered in our analysis. Coordinates for the Center Of Gravity (COG) of the peak activation are given in mm space for X, Y and Z coordinates and size was determined based on the number of vertices in each cluster. The name of each area was determined by entering the coordinates into Neurosynth and using the top gyri/sulci name. R = right hemisphere and L = left hemisphere. DLPFC = Dorsal lateral prefrontal cortex, PCC = Posterior Cingulate Cortex, SMA = Somatamotor area.
\*clusters that replicated in the sample split half replication.

Our method also creates an environmental association map. If genetic and environmental association are in the same direction, it is consistent with an explanation of causality(39), though not sufficient to establish a causal relationship. Environmental associations were not consistently in the same direction of effect as the genetic clusters that were discovered (see Supplemental Figure S5). Thus, from the environmental map analysis and the bivariate Cholesky decomposition of ICU and CESD, we concluded these areas are likely biomarkers that reflect genetic vulnerability to CU and CESD and implicate a shared genetic liability.

### What Genetic Pathways are Implicated?

We used follow-up analyses to gain insight into potential mechanisms involved in this genetic vulnerability. Results of clustering of Psychiatric Genomics Consortium depression-related genes are shown in Figure 4. We found three clusters: overexpressed, mixed expression, and under-expressed (genes listed in axis of Figure 4). The overexpressed cluster showed significant enrichment for genes in “Depolarization of the Presynaptic Terminal Triggers the Opening of Calcium Channels_Homo sapiens_R-HSA-112308” pathway (FDR corrected *p*=.03). No other pathways were significant across any of our clusters after FDR correction.

**Figure 4.**
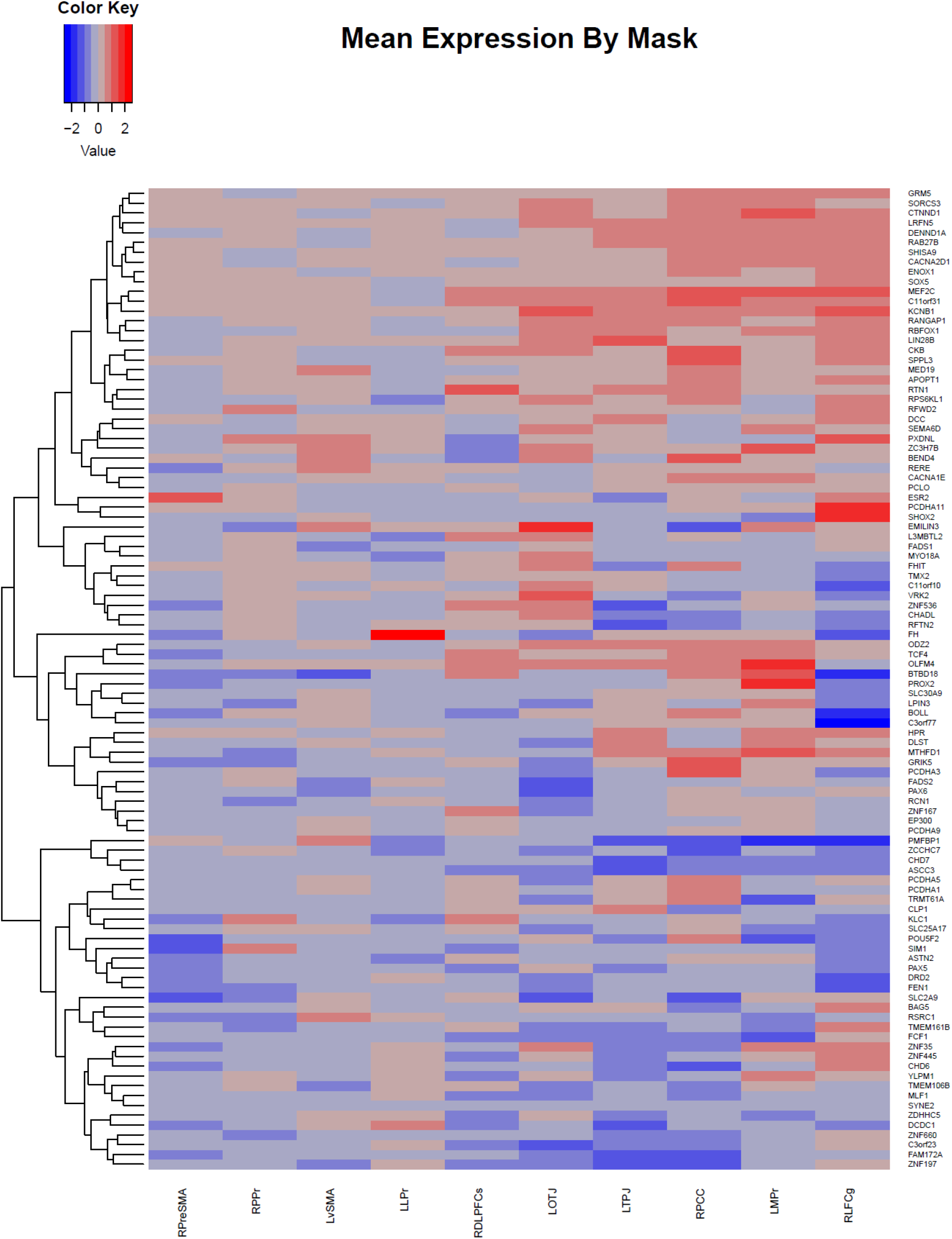
Hierarchical clustering of expression patterns of depression genes in derived clusters. Color scale is the z-score for the degree of expression of that gene in the derived area mask compared to the whole cortex. Depression genes were obtained from the Psychiatric Genomics Consortium GWAS gene-burden tests, excluding genes from the major histocompatibility complex region(1). Gene expression values were recovered from Neurosynth-gene, which processed data from the Allen Brain Atlas, Human Brain Atlas. R = right hemisphere clusters; L = left hemisphere clusters. RDLPFCs = right dorsal lateral prefrontal cortex sulci, RLFCg = Right Lateral frontal gyri, LLPr = Left Lateral Precuneus, LMPr = Left Medial Precuneus, LOTJ = Left Occipital Temporal Junction, RPreSMA = Right Pre-Somatosensory Area, RPCC = Right Posterior Cingulate Cortex, LvSMA = Left Ventral somatosensory, LTPJ = Left temporoparietal Junction, and RPPr= Right Posterior Precuneus.

### What Likely Cognitive/Behavioral Pathways are Involved?

To identify likely cognitive/behavioral mechanisms reflecting this vulnerability, we conducted a meta-analytic term search using Neurosynth. Supplemental Tables S4 and S5 show the 25 most positively associated function terms from Neurosynth for each genetic overlap cluster from the full sample (in some cases, fewer than 25 terms were positively associated). The top repeated behavioral terms were “Theory of Mind”, “inhibit”, and “pain” across all regions (using a wildcard* for different forms of the same word and spelling out acronyms).

We projected our genetic derived clusters onto the Yeo 7-network parcellation, a popular, low-dimensionality parcellation derived from a clustering analysis of resting state data from 1000 participants(35). Supplemental Table S6 reports the results of this analyses. The default network was the most common network (4 areas); all but one positively associated cluster from our genetic analysis fell in this network, in line with past research that implicated default network functions to depression (40). All but two areas (8 of 10 positively and negatively associated areas) fell in networks with higher-level cognitive functions (i.e., default mode, ventral and dorsal attention, and frontal networks).

## Discussion

By directly estimating brain areas genetically associated with depression and CU we found (1) the association between CU and depressive symptoms was entirely genetic in origin. (2) Genetic influences on thicker cortex in right DLFPC sulci, the right PreSMA, left medial and lateral precuneus, OTJ, and TPJ were associated with both traits, and genetic influences on thinner cortex in the right ventral PCC, right medial precuneus, right DLPFC gyrus, and left somatosensory cortex were associated with both traits. (3) Likely molecular pathways are influencing calcium channel depolarization. (4) Likely associated behaviors are “theory of mind”, “inhibit”, and “pain”. (5) Connectivity is associated with default-mode and higher-level cognitive systems. Figure 5 links our results across different methods to the RDoC social dimensions matrix. We discuss the implications of these findings below.

**Figure 5.**
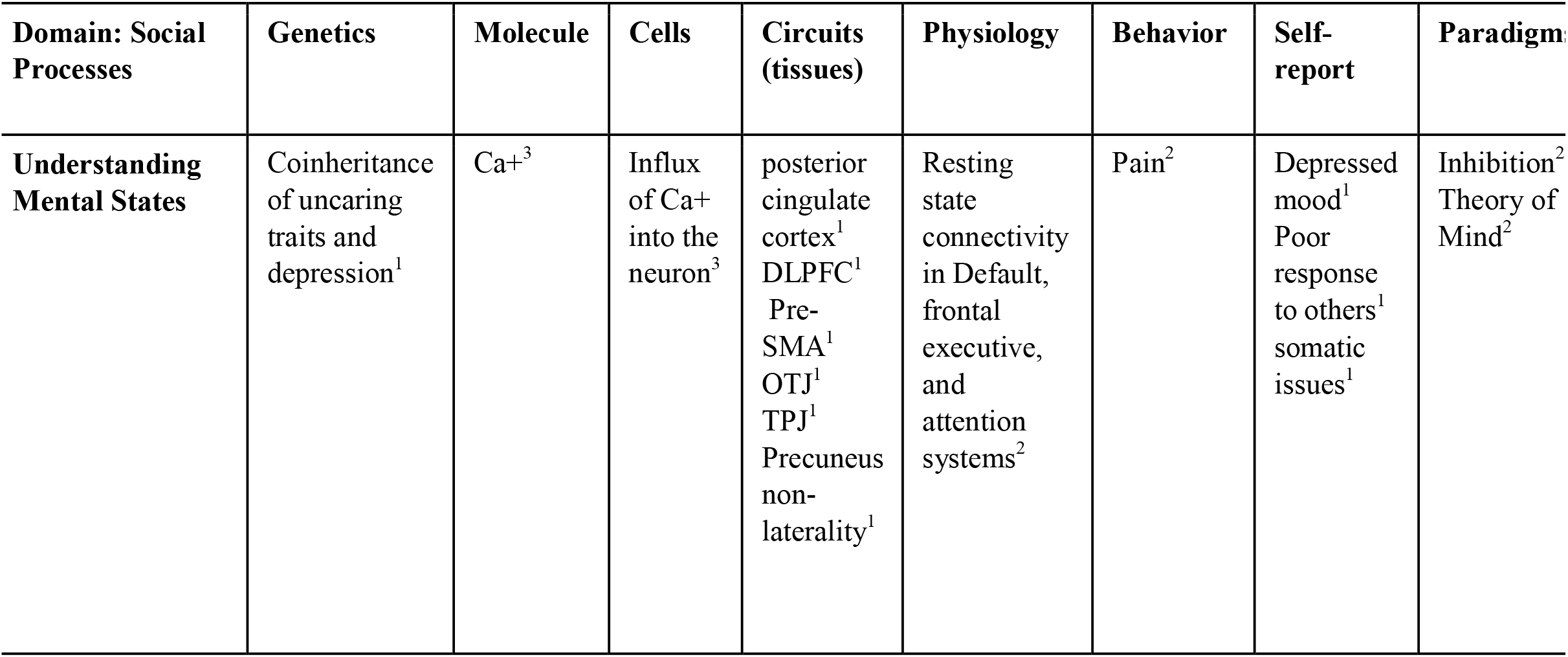
We used the RDoC “Social Processing: Understanding Mental States” domain dimension matrix to organize our results across different levels of biology and literature. DLPFC=Dorsal Lateral Prefrontal Cortex, Pre-SMA=Pre-Somatosensory Area, OTJ=Occipital Temporal Junction, TPJ = temporoparietal Junction, Ca+=Calcium, positive charge. ^1^Results were estimated directly in this study. ^2^Results were found using MNI coordinates that overlap spatially with those found in the fMRI literature, including the Yeo 7 networks and Neurosynth meta-analytic database. ^3^Results use the Allen Brain Atlas to visualize expression of PGC MDD associated genes.

### Advantages of Brain Mapping Approach

We are the first to directly estimate the cortical pattern that represents genetic vulnerability to a psychiatric disorder. Importantly, this approach is not limited to known associations (i.e., brain regions that are not phenotypically associated with depression can reflect genetic vulnerability due to sampling error and environmental associations), and can account for the architecture of genetic effects on brain structure(41). Further, our approach allows for expansion of hypotheses in genetic association studies by integrating MRI atlas-based approaches to contextualize the genetic association patterns and implicate molecular pathways and brain functions.

In this case, we focused on the vulnerability for CU in depression, chosen due to its importance in depression severity(42) and integration with RDoC domains(2). Reassuringly, this approach converges on several areas previously associated with depression and social processing literature(17, 22), which means past neuroimaging studies of these behaviors may be driven by genetics. However, cortical thickness associations with depression in the temporoparietal and temporo-occipital junctions, key social processing areas, were novel. Finally, we identified likely mechanisms for follow-up analyses using Bayesian meta-analysis, such as theory of mind and inhibition, that are likely targets for behavioral intervention.

This vulnerability reflects an expanded cognitive network. We found clusters specific to the posterior ventral cingulate cortex and DLFPC, which show broad connectivity patterns (functional and anatomical) between limbic/emotional systems and the association cortex(45, 46, 47). Further, almost all clusters were in higher-order cognitive systems. At the molecular level, we implicated positively charged calcium channels. Further informatic analysis implicated theory of mind, meaning talk therapy may be an effective intervention target, with adaptions of interpersonal mindfulness showing efficacy for depression(46).

### Limitations

There are limitations to our approach. First, the sample is tightly matched on age (at around age 28). While this protects against both linear and non-linear confounding by age and developmental effects, results may not generalize to young or old age cohorts. Second, we did not have enough power to explore interactions with sex. Although sex interaction may be a factor in genetic depressive symptomology, there is still a genetic correlation between males and females(14). Additionally, informatic analyses focused on overlap based on spatial coordinates. While inclusion of results from neuroinformatic tools is more expansive, we did not explicitly model the patterns of association between RNA transcripts, inferred behaviors, etc.

## Conclusion

We directly mapped genetic vulnerability to CU and depressive symptoms on the brain. We found common genetic variance in CU and depressive symptoms was associated with higher-order cognitive areas and functions. As the genetic vulnerability to psychiatric disorders is discovered, the use of high-resolution cortical methods will be invaluable in contextualizing the patterns of genetic effects.

## Acknowledgments

This research was supported by NIH grants MH063207, AG046938, HD010333, and MH016880.

## References

1. Wray NR, Ripke S, Mattheisen M, Trzaskowski M, Byrne EM, Abdellaoui A, et al. (2018): Genome-wide association analyses identify 44 risk variants and refine the genetic architecture of major depression. Nat Genet. 50: 668–681.

2. Insel T, Cuthbert B, Garvey M, Heinssen R, Pine DS, Quinn K, et al. (2010): Research Domain Criteria (RDoC): Toward a new classification famework for research on mental disorders. Am J Psychiatry. 167: 748–751.

3. Fekadu A, Wooderson SC, Markopoulo K, Donaldson C, Papadopoulos A, Cleare AJ (2009): What happens to patients with treatment-resistant depression? A systematic review of medium to long term outcome studies. J Affect Disord. 116: 4–11.

4. Schreiter S, Pijnenborg GHM, aan het Rot M (2013): Empathy in adults with clinical or subclinical depressive symptoms. J Affect Disord. 150: 1–16.

5. Wolkenstein L, Schönenberg M, Schirm E, Hautzinger M (2011): I can see what you feel, but I can’t deal with it: Impaired theory of mind in depression. J Affect Disord. 132: 104–111.

6. Cusi AM, MacQueen GM, Spreng RN, McKinnon MC (2011): Altered empathic responding in major depressive disorder: Relation to symptom severity, illness burden, and psychosocial outcome. Psychiatry Res. 188: 231–236.

7. Ladegaard N, Larsen ER, Videbech P, Lysaker PH (2014): Higher-order social cognition in first-episode major depression. Psychiatry Res. 216: 37–43.

8. Centifanti LCM, Meins E, Fernyhough C (2016): Callous-unemotional traits and impulsivity: distinct longitudinal relations with mind-mindedness and understanding of others. J Child Psychol Psychiatry 57: 84–92.

9. Frick PJ, Ellis M (1999): Callous-unemotional traits and subtypes of conduct disorder. Clin Child Fam Psychol Rev. 2: 149–168.

10. Kimonis ER, Frick PJ, Skeem JL, Marsee MA, Cruise K, Munoz LC, et al. (2008): Assessing callous-unemotional traits in adolescent offenders: Validation of the Inventory of Callous-Unemotional Traits. Int J Law Psychiatry. 31: 241–252.

11. Byrd AL, Kahn RE, Pardini DA (2013): A Validation of the Inventory of Callous-Unemotional Traits in a community sample of young adult males. J Psychopathol Behav Assess. 35. doi: 10.1007/s10862-012-9315-4.

12. Lamers F, de Jonge P, Nolen WA, Smit JH, Zitman FG, Beekman ATF, Penninx BWJH (2010): Identifying depressive subtypes in a large cohort study. J Clin Psychiatry. 71: 1582–1589.

13. Carney DM, Moroney E, Machlin L, Hahn S, Savage JE, Lee M, et al. (2016): The Twin Study of Negative Valence Emotional Constructs. Twin Res Hum Genet. 19: 456–64.

14. Sullivan PF, Neale MC, Kendler KS (2000): Genetic epidemiology of major depression: review and meta-analysis. Am J Psychiatry. 157: 1552–1562.

15. Levinson DF, Mostafavi S, Milaneschi Y, Rivera M, Ripke S, Wray NR, Sullivan PF (2014): Genetic studies of major depressive disorder: why are there no genome-wide association study findings and what can we do about it? Biol Psychiatry. 76: 510–2.

16. Sunkin SM, Ng L, Lau C, Dolbeare T, Gilbert TL, Thompson CL, et al. (2013): Allen Brain Atlas: an integrated spatio-temporal portal for exploring the central nervous system. Nucleic Acids Res. 41: D996–D1008.

17. Schmaal L, Hibar DP, Sämann PG, Hall GB, Baune BT, Jahanshad N, et al. (2017): Cortical abnormalities in adults and adolescents with major depression based on brain scans from 20 cohorts worldwide in the ENIGMA Major Depressive Disorder Working Group. Mol Psychiatry. 22: 900–909.

18. Uddin LQ (2015): Salience processing and insular cortical function and dysfunction. Nat Rev Neurosci. 16: 55–61.

19. Andrews-Hanna JR, Reidler JS, Sepulcre J, Poulin R, Buckner RL (2010): Functional-anatomic fractionation of the brain’s default network. Neuron. 65: 550–562.

20. The salience network is responsible for switching between the default mode network and the central executive network: Replication from DCM (2014): Neuroimage. 99: 180–190.

21. Fischl B, Sereno MI, Tootell RBH, Dale AM (1999): High-resolution intersubject averaging and a coordinate system for the cortical surface. Hum Brain Mapp. 8: 272–284.

22. Bora E, Fornito A, Pantelis C, Yücel M (2012): Gray matter abnormalities in Major Depressive Disorder: a meta-analysis of voxel based morphometry studies. J Affect Disord. 138: 9–18.

23. De Brito SA, Mechelli A, Wilke M, Laurens KR, Jones AP, Barker GJ, et al. (2009): Size matters: Increased grey matter in boys with conduct problems and callous-unemotional traits. Brain. 132: 843–852.

24. Wagner G, Koch K, Schachtzabel C, Schultz CC, Sauer H, Schlösser RG (2011): Structural brain alterations in patients with major depressive disorder and high risk for suicide: Evidence for a distinct neurobiological entity? Neuroimage. 54: 1607–1614.

25. Amunts K, Schleicher A, Bürgel U, Mohlberg H, Uylings HB, Zilles K (1999): Broca’s region revisited: cytoarchitecture and intersubject variability. J Comp Neurol. 412: 319–41.

26. Eyler LT, Chen C-H, Panizzon MS, Fennema-Notestine C, Neale MC, Jak A, et al (2012): A comparison of heritability maps of cortical surface area and thickness and the influence of adjustment for whole brain measures: a magnetic resonance imaging twin study. Twin Res Hum Genet. 15: 304–14.

27. Eaton, W. W., Smith, C., Ybarra, M., Muntaner, C., & Tien A (2004): Center for Epidemiologic studies depression scale: Review and revision (CESD and CESD-R). (M. E. Maruish, editor). Mahwah, NJ: Lawrence Erlbaum Associates. Retrieved July 26, 2017, from http://psycnet.apa.org/record/2004-14941-011.

28. Friedman NP, du Pont A, Corley RP, Hewitt JK (2018): Longitudinal relations between depressive symptoms and executive functions from adolescence to early adulthood: A twin study. Clin Psychol Sci. 6: 543–560.

29. Byrd AL, Kahn RE, Pardini DA (2013): A validation of the Inventory of Callous-Unemotional Traits in a community sample of young adult males. J Psychopathol Behav Assess. 35. doi: 10.1007/s10862-012-9315-4.

30. Hagler DJ, Saygin AP, Sereno MI, Sereno MI (2006): Smoothing and cluster thresholding for cortical surface-based group analysis of fMRI data. Neuroimage. 33: 1093–103.

31. Neale MC, Cardon LR (1992): Methodology for Genetic Studies of Twins and Families. Dordrecht: Springer Netherlands. doi: 10.1007/978-94-015-8018-2.

32. Fischl B (2012): FreeSurfer. Neuroimage. 62: 774–781.

33. Friston KJ, Penny WD, Glaser DE (2005): Conjunction revisited. Neuroimage. 25: 661–667.

34. Yarkoni T, Poldrack RA, Nichols TE, Van Essen DC, Wager TD (2011): Large-scale automated synthesis of human functional neuroimaging data. Nat Methods. 8: 665–670.

35. Yeo BTT, Krienen FM, Sepulcre J, Sabuncu MR, Lashkari D, Hollinshead M, et al. (2011): The organization of the human cerebral cortex estimated by intrinsic functional connectivity. J Neurophysiol. 106: 1125–65.

36. Fox AS, Chang LJ, Gorgolewski KJ, Yarkoni T (2014): Bridging psychology and genetics using large-scale spatial analysis of neuroimaging and neurogenetic data. doi.org. 012310.

37. Crum WR (2011): Magnetic resonance brain image processing and arithmetic with FSL. Humana Press, pp 109–126.

38. Joshi-Tope G, Gillespie M, Vastrik I, D’Eustachio P, Schmidt E, de Bono B, et al. (2004): Reactome: a knowledgebase of biological pathways. Nucleic Acids Res. 33: D428–D432.

39. Heath AC, Kessler RC, Neale MC, Hewitt JK, Eaves LJ, Kendler KS (1993): Testing hypotheses about direction of causation using cross-sectional family data. Behav Genet. 23: 29–50.

40. Kaiser RH, Andrews-Hanna JR, Wager TD, Pizzagalli DA, Rj D, J J (2015): Large-scale network dysfunction in major depressive disorder. JAMA Psychiatry. 72: 603.

41. Rimol LM, Panizzon MS, Fennema-Notestine C, Eyler LT, Fischl B, Franz CE, et al. (2010): Cortical thickness is influenced by regionally specific genetic factors. Biol Psychiatry. 67: 493–499.

42. Szanto K, Dombrovski AY, Sahakian BJ, Mulsant BH, Houck PR, Reynolds CF, Clark L (2012): Social emotion recognition, social functioning, and attempted suicide in late-life depression. Am J Geriatr Psychiatry. 20: 257–265.

43. Leech R, Sharp DJ (2014): The role of the posterior cingulate cortex in cognition and disease. Brain. 137: 12–32.

44. Pizzagalli DA (2010): Frontocingulate dysfunction in depression: Toward biomarkers of treatment response. Neuropsychopharmacol Publ online 22 Sept 2010; doi 1138/npp201166.36: 183.

45. Goldapple K, Segal Z, Garson C, Lau M, Bieling P, Kennedy S, et al. (2004): Modulation of cortical-limbic pathways in major depression. Arch Gen Psychiatry. 61: 34.

46. Harley R, Sprich S, Safren S, Jacobo M, Fava M (2008): Adaptation of dialectical behavior therapy skills training group for treatment-resistant depression. J Nerv Ment Dis. 196: 136–143.

