## Supplemental Information 1 for "Whole-Cortex Mapping of Common Genetic Influences on Depression and Social Deficits"

Supplement

Index

I. Online methods
   a. Sample acquisition
   b. MRI and questionnaire procedures
II. Supplemental results
    a. Tables
       1. Descriptive statistics for CESD, ICU, and ICU subscales
       2. Full ACE model fitting results for CESD and ICU
       3. Genetic associations between ICU subscales and CESD.
       4. Full results of Neurosynth term search for each left hemisphere cluster
       5. Full results of Neurosynth term search for each right hemisphere cluster
       6. Spatial overlap between each cluster and Yeo 7 networks
    b. Figures
       1. P-factor model to demonstrate specificity of association
       2. Map of C effects across the cortex
       3. Plot of elbow method results for K-means clustering
       4. Overlap from split-half replicability analysis
       5. Map of environmental pattern and overlap with discover clusters
       6. Standard (phenotypic) brain map results for ICU and CESD

Online Methods

A. *Sample and Acquisition*

The Longitudinal Twin Study (LTS) is composed of adult individuals that were ascertained through the Twin Infant Project in 1984, and the MacArthur Longitudinal Twin Study in 1986. The Colorado Department of Health solicited participation in the registry using birth records of all families in which (1) both twins survived, (2) were healthy and (3) were within ~3 hours driving time of CU-Boulder. The sample is relatively representative of twin births for the time considering these criteria (Rhea et al., 2006). We tested all eligible individuals (those who consented and could safely enter the scanning environment), regardless of handedness, medication status, head injury history, or substance use history. We asked subjects to contribute a urine and saliva sample to assess current levels of various substances. About 30% of the sample no longer lives locally. These subjects traveled to Colorado for the imaging sessions.

B. *MRI and Questionnaire Procedures*

After completing informed consent, participants completed screening measures and the Center for Epidemiological Studies Depression scale (CESD), and were informed about scanning procedures. The Inventory of Callous and Unemotional traits (ICU) was collected online prior to the scanning session.

Cortical thickness was calculated with the Freesurfer analysis suite (<http://surfer.nmr/mgh.harvard.edu>)(1). T1-wieghted images were extracted using watershed/surface deformation procedure(2), surface deformation along intensity gradients to optimally differentiate gray matter, white matter and cerebral spinal fluid boundaries(3), and tessellation of gray/white matter boundary(4). The resulting surfaces were registered to standard spherical brain template(5, 6), and then used to compute a range of surface-based measurements. Finally, vertices were smoothed at 10mm across the cortex full-width-at-half-maximum (FWHM) isotropic kernel(7).

Supplemental Table S1. *Descriptive Statistics for CESD, ICU, and ICU Subscales*

| **Scale** | **Mean** | **SD** | **Min** | **Max** | **Cronbach's Alpha** | **Shapiro-Wilk for Raw Scores** | **Shapiro-Wilk for**  **Square Root Transformed Scores** |
| --- | --- | --- | --- | --- | --- | --- | --- |
| CESD | 8.786 | 8.477 | 0 | 50.000 | 0.902 | 0.834 | 0.979 |
| ICU total | 15.595 | 6.389 | 1 | 38.000 | 0.771 | 0.984 | 0.992 |
| ICU callous | 2.118 | 2.345 | 0 | 18.000 | 0.595 | 0.81 | 0.915 |
| ICU unemotional | 6.180 | 2.959 | 0 | 15.000 | 0.814 | 0.974 | 0.912 |
| ICU uncaring | 5.453 | 3.326 | 0 | 17.000 | 0.736 | 0.961 | 0.973 |

*Note.* Descriptive statistics for the the Center for Epidemiological Studies (CESD) total scale and the Inventory of Callous and Unemotional traits (ICU) total scale and subscales.

Supplemental Table S2. *Univariate Twin Models for Behavioral Measures*

|  | **Model Fit** | | | | | | **Standardized Paths** | | | |
| --- | --- | --- | --- | --- | --- | --- | --- | --- | --- | --- |
| **Model** | **χ^2^** | **df** | ***p*** | **AIC** | **BIC** | **RMSEA** | **A** | **C** | **D** | **E** |
| CESD |  |  |  |  |  |  |  |  |  |  |
| ACE | 11.164 | 6 | 0.0834 | 1745.565 | 1760.230 | 0.077 | 0.597 | 0.000 | -- | 0.802 |
| ADE | 10.509 | 6 | 0.1048 | 1744.910 | 1759.576 | 0.072 | 0.002 | -- | 0.610 | 0.793 |
| **AE** | **11.164** | **7** | **0.1316** | **1743.565** | **1754.564** | **0.064** | **0.597** | **--** | **--** | **0.802** |
| CE | 33.427 | 7 | <.0001 | 1765.828 | 1776.827 | 0.162 | -- | 0.707 | -- | 0.707 |
| DE | 10.509 | 7 | 0.1615 | 1742.910 | 1753.909 | 0.059 | -- | -- | 0.610 | 0.793 |
| E | 33.427 | 8 | 0.0001 | 1763.828 | 1771.161 | 0.148 | -- | -- | -- | 1.000 |
| ICU |  |  |  |  |  |  |  |  |  |  |
| ACE | 10.722 | 6 | 0.0974 | 1189.329 | 1203.939 | 0.074 | 0.574 | 0.244 | NA | 0.782 |
| ADE | 10.793 | 6 | 0.0950 | 1189.400 | 1204.010 | 0.075 | 0.627 | -- | 0.000 | 0.779 |
| **AE** | **10.793** | **7** | **0.1479** | **1187.400** | **1198.358** | **0.062** | **0.627** | **--** | **--** | **0.779** |
| CE | 12.597 | 7 | 0.0826 | 1189.204 | 1200.161 | 0.075 | -- | 0.575 | 0.818 | 0.818 |
| DE | 12.018 | 7 | 0.1000 | 1188.625 | 1199.582 | 0.071 | -- | -- | 0.631 | 0.776 |
| E | 38.447 | 8 | <.0001 | 1213.054 | 1220.359 | 0.163 | -- | -- | -- | 1.000 |

*Note.* Model fit for univariate models of Center for Epidemiological Studies-Depression scale (CESD) and the Inventory of Callous and Unemotional traits (ICU) total score. The CESD and ICU were residualized on mean thickness and sex. For each model, we tested what combination of A (Additive genetic), D (Dominance genetic), C (Common environment), or E (nonshared Environment) best fit each scale. Dashes indicates that the parameter was not estimated in that particular model. We used χ^2^ difference testing, AIC, BIC and Root Mean Square Error of Approximation (RMSEA) as standards for model comparison. The preferred model is indicated in bold-face type. Although the AE and DE models fit similarly, the AE model was preferred, because D without A variance is biologically implausible. *N*= 285 same-sex twin pairs (142 monozygotic [MZ] and 143 dizygotic [DZ]). This sample was larger than that in the main analysis because it included twin pairs that were missing brain data but had behavioral assessments.

Supplemental Table S3. *Genetic Association Between CESD and Each ICU Subscale*

| **Var1** | **Path1A** | ***rG*** | **Gr** | ***Pvalue*** |
| --- | --- | --- | --- | --- |
| Callousness | 0.456374 | 0.029352 | 0.00763 | 0.215 |
| Uncaring | 0.520091 | 0.491772 | 0.151531 | 0.02 |
| Unemotional | 0.790123 | 0.08316 | 0.038975 | 0.502 |
| Total Scale | 0.585954 | 0.351804 | 0.12266 | 0.036 |

*Note.* Center for Epidemiological Studies Depression scales (CESD) and Inventory of Callous and Unemotional traits (ICU) were residualized on sex and mean thickness. *r*G represents the genetic correlation and bivariate heritability represents the phenotypic correlation predicted by genetic covariance. All estimates were derived from the standard bivariate Cholesky decomposition. The A path represents each subscale's standardized A estimate; squaring this value yields the heritability. The *p*-value for the genetic associations was estimated by a 1-df chi-square difference test of the Cholesky cross path.

Supplemental Table S4. *Meta-analytic Terms for Right Hemisphere Clusters*

| **R-PreSMA** | | **R-Precuneus** | | **R-PCC** | | **R-Frontal Lateral Gyri** | | **R-Frontal Sulci** | |
| --- | --- | --- | --- | --- | --- | --- | --- | --- | --- |
| **Term** | ***R*** | **Term** | ***r*** | **Term** | ***r*** | **Term** | **r** | **Term** | ***r*** |
| conflict | 0.164 | causality | 0.119 | inhibitory | 0.143 | None |  | reducing | 0.3 |
| distractors | 0.145 | precuneus posterior | 0.106 | Inhibit | 0.119 |  |  | relied | 0.115 |
| cortex anterior | 0.077 | experimentally | 0.101 | abnormality | 0.108 |  |  | middle cingulate | 0.085 |
| orienting | 0.07 | precuneus | 0.096 | prefrontal cortices | 0.089 |  |  | imagine | 0.067 |
| cortex acc | 0.056 | centered | 0.09 | chronic pain | 0.075 |  |  | amygdala anterior | 0.039 |
| acc | 0.053 | cortex precuneus | 0.087 | nervous | 0.074 |  |  | prefrontal cortex | 0.02 |
| dorsolateral prefrontal | 0.036 | theory | 0.084 | posterior medial | 0.037 |  |  | cingulate | 0.018 |
| anterior cingulate | 0.035 | deactivations | 0.082 | cingulate | 0.023 |  |  | prefrontal | 0.018 |
| task | 0.035 | midline | 0.082 | anterior insula | 0.01 |  |  |  |  |
| anterior | 0.03 | spontaneous | 0.082 | brainstem | 0.009 |  |  |  |  |
| cingulate cortex | 0.029 | thoughts | 0.08 | midbrain | 0.008 |  |  |  |  |
| anterior insula | 0.025 | mode | 0.077 | pain | 0.007 |  |  |  |  |
| cingulate | 0.025 | posterior cingulate | 0.077 |  |  |  |  |  |  |
| supplementary | 0.023 | Pcc | 0.075 |  |  |  |  |  |  |
| supplementary motor | 0.023 | default mode | 0.073 |  |  |  |  |  |  |
| parietal cortex | 0.022 | mode network | 0.072 |  |  |  |  |  |  |
| execution | 0.02 | connectivity networks | 0.071 |  |  |  |  |  |  |
| basal ganglia | 0.019 | default | 0.07 |  |  |  |  |  |  |
| ganglia | 0.019 | Self | 0.068 |  |  |  |  |  |  |
| prefrontal | 0.018 | preserved | 0.066 |  |  |  |  |  |  |
| motor | 0.009 | striking | 0.066 |  |  |  |  |  |  |
|  |  | mental | 0.065 |  |  |  |  |  |  |
|  |  | theory mind | 0.065 |  |  |  |  |  |  |
|  |  | deactivation | 0.064 |  |  |  |  |  |  |
|  |  | independent component | 0.064 |  |  |  |  |  |  |

*Note.* Top associated terms for a Neurosynth decoder analysis for each right hemisphere overlay cluster. Decoder analysis uses a meta-analytic method to find terms that appear frequently in papers that report effects at the coordinates input into the analyses. Each cluster was estimated as a mask and then put through decoder analysis separately. Each term was put into a single vector (excluding repeating terms for each mask) and the most common words across all areas were reported in the main text, using a wildcard* to account for like words.

Supplemental Table S5. *Meta-analytic Terms for Left Hemisphere Clusters*

| **L-vSMA** | | **L-PTJ** | | **L-OTJ** | | **L-Precuneus** | |
| --- | --- | --- | --- | --- | --- | --- | --- |
| **Term** | ***R*** | **Term** | ***r*** | **Term** | ***r*** | **Term** | ***r*** |
| receiving | 0.221 | having | 0.333 | picture | 0.295 | personal | 0.1 |
| practice | 0.18 | indirect | 0.284 | researchers | 0.109 | sparse | 0.069 |
| facilitated | 0.13 | trained | 0.138 | videos | 0.1 | thinking | 0.065 |
| touch | 0.117 | failure | 0.134 | convergence | 0.094 | cortex posterior | 0.061 |
| sii | 0.09 | temporo parietal | 0.126 | actively | 0.091 | maintaining | 0.061 |
| tactile | 0.09 | inhibit | 0.123 | monkey | 0.086 | precuneus | 0.051 |
| si | 0.088 | temporo | 0.113 | extrastriate | 0.075 | cingulate cortices | 0.043 |
| sensory | 0.079 | recording | 0.081 | selectivity | 0.075 | states | 0.043 |
| induced | 0.075 | access | 0.08 | pictures | 0.07 | nervous | 0.037 |
| secondary somatosensory | 0.075 | online | 0.076 | scene | 0.062 | dorsomedial prefrontal | 0.032 |
| s1 | 0.074 | response inhibition | 0.076 | motion | 0.061 | mental states | 0.025 |
| somatosensory cortex | 0.072 | valid | 0.068 | primary visual | 0.061 | mind tom | 0.022 |
| stimulation | 0.068 | mind | 0.054 | consecutive | 0.06 | posterior cingulate | 0.022 |
| somatosensory | 0.067 | reappraisal | 0.054 | middle occipital | 0.052 | tom | 0.022 |
| primary secondary | 0.065 | error | 0.052 | mt | 0.052 | autobiographical memory | 0.018 |
| lobule ipl | 0.064 | links | 0.046 | extrastriate visual | 0.051 | default mode | 0.018 |
| Painful | 0.061 | endogenous | 0.044 | plus | 0.051 | mode | 0.017 |
| primary somatosensory | 0.06 | theory mind | 0.042 | moving | 0.048 | mode network | 0.016 |
| discriminative | 0.059 | mind tom | 0.04 | v1 | 0.045 | theory mind | 0.016 |
| sparse | 0.058 | successful | 0.039 | direction | 0.044 | episodic | 0.015 |
| motor control | 0.057 | group healthy | 0.037 | occipital | 0.044 | default | 0.014 |
| matrix | 0.056 | situation | 0.036 | body | 0.043 | autobiographical | 0.012 |
| motor task | 0.056 | actively | 0.032 | parieto | 0.043 | cingulate | 0.011 |
| secondary | 0.053 | mtg | 0.032 | social cognition | 0.043 |  |  |
| pain | 0.052 | tom | 0.032 | vision | 0.041 |  |  |

*Note.* Top associated terms for a Neurosynth decoder analysis for each left hemisphere overlay cluster. Decoder analysis uses a meta-analytic method to find terms that appear frequently in papers that report effects at the coordinates input into the analyses. Each cluster was estimated as a mask and then put through decoder analysis separately. Each term was put into a single vector (excluding repeating terms for each mask) and the most common words across all areas were reported in the main text, using a wildcard* to account for like words.

Supplemental Table S6. *Spatial Overlap of Overlap Clusters with the Yeo 7 Networks*

| **Overlap Cluster** | **Network In Yeo 7** | **Direction** |
| --- | --- | --- |
| L-OTJ | Visual | + |
| L-TPJ | Default | + |
| L-vSMA | Somatamotor | - |
| L-Medial Precuneus | Dorsal Attention | + |
| L-Lateral Precuneus | Default | + |
| R-DLPFC sulci | Default | + |
| R-DLPFC Gyri | Frontal | - |
| R-Posterior Cingulate | Frontal | - |
| R-PreSMA | Ventral Attention | + |
| R-Medial Precuneus | Default | - |

*Note.* Each cluster fell into only one network. The direction of effect in the original anatomical genetic brain map is shown to the right. Overlap is based on overlap of spatial coordinates, and not a statistical test of association. L- = left hemisphere and R- = right hemisphere. OTJ=Occipital Temporal Junction, TPJ = Parietal Temporal Junction, vSMA=ventral somatosensory Motor area, DLPFC = Dorsal Lateral Prefrontal Cortex, PreSMA = Pre-Somatosensory area.

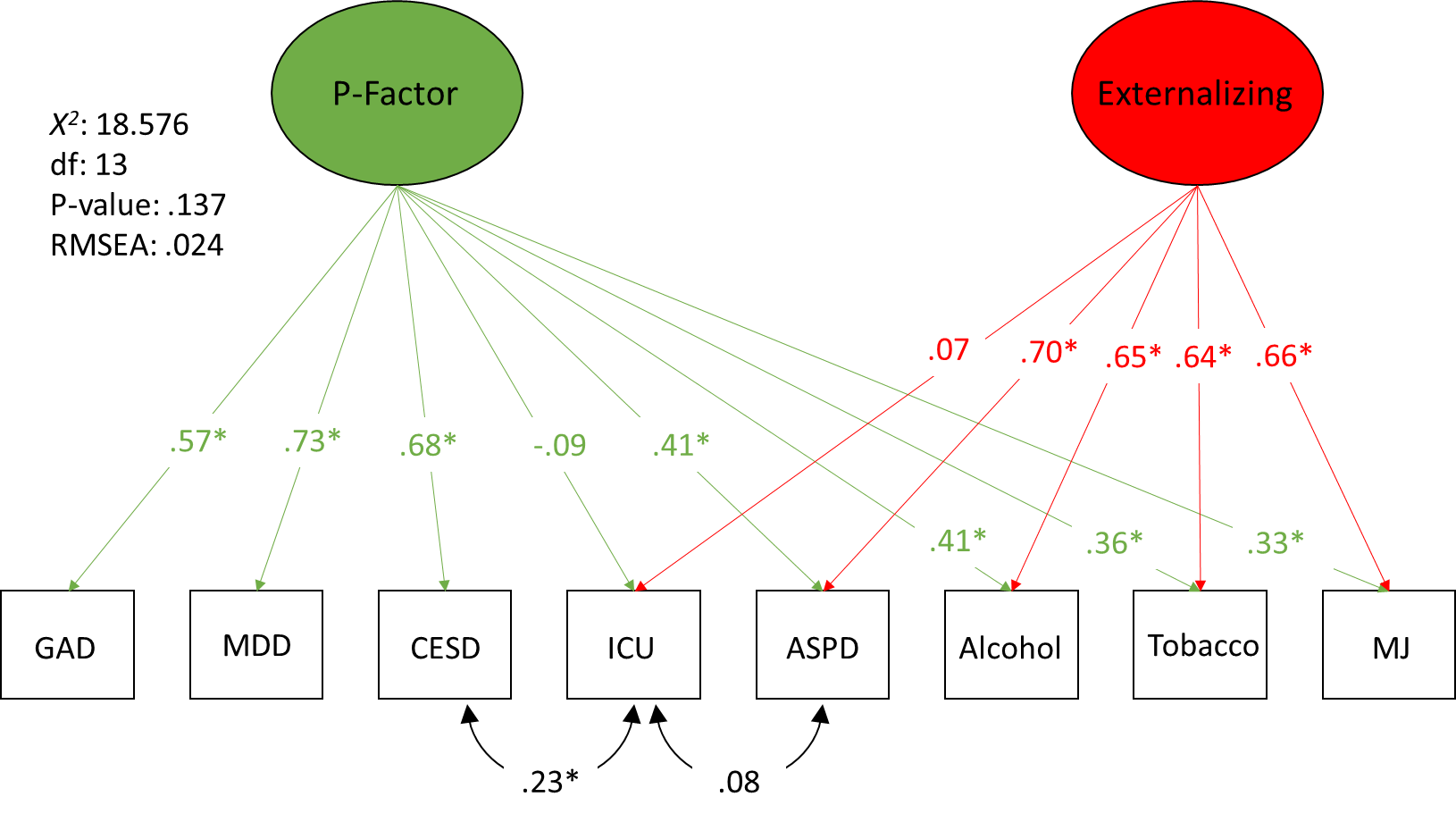

*Supplementary Figure S1.* P-factor model. The indicators for Generalized Anxiety Disorder (GAD), Major Depressive Disorder (MDD), Antisocial Personality Disorder (ASPD), Alcohol addiction (Alcohol), Tobacco addiction (Tobacco) and Marijuana addiction (MJ) were lifetime DSM-4 diagnosis, based on structured clinical interviews at age 23(8). The diagnosis variables were coded as 0 for no symptoms, 1 for symptoms but no diagnosis, and 2 for diagnosis. We treated these variables as ordinal with a threshold model, estimated with the weighted least squares, means and variance adjusted (WLSMV) estimator. Fit statistics for this model are shown to the left. We tested for a specific association (above and beyond the P-factor) between the CESD and ICU by adding a residual correlation. We also added a residual correlation between the ICU and ASPD, as these variables have been associated in the past. The model includes a P-factor and Externalizing factor only because the two measures that would normally load on an Internalizing factor left the model empirically underidentified when allowed to load on their own factor due to low factor loadings. **p* < .05.

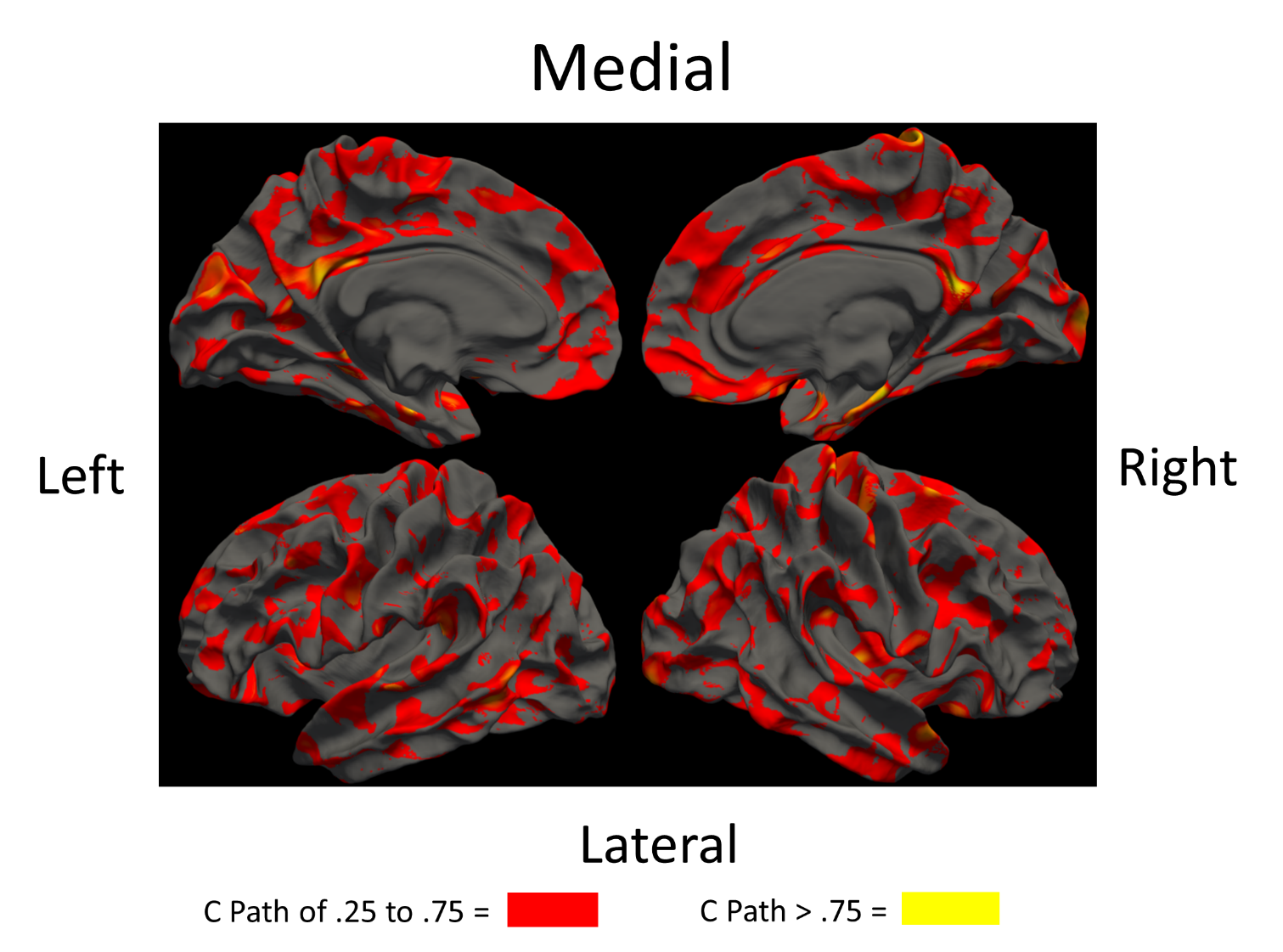

*Supplemental Figure S2.* Map of the shared environmental (C) effects for thickness across the whole cortex, thresholded by path estimate. C effects range from explaining 5 to over 50% of the variance across the brain. These estimates were taken from the same map as the Cholesky decomposition for the CESD to demonstrate the substantial C those models found across the cortex.

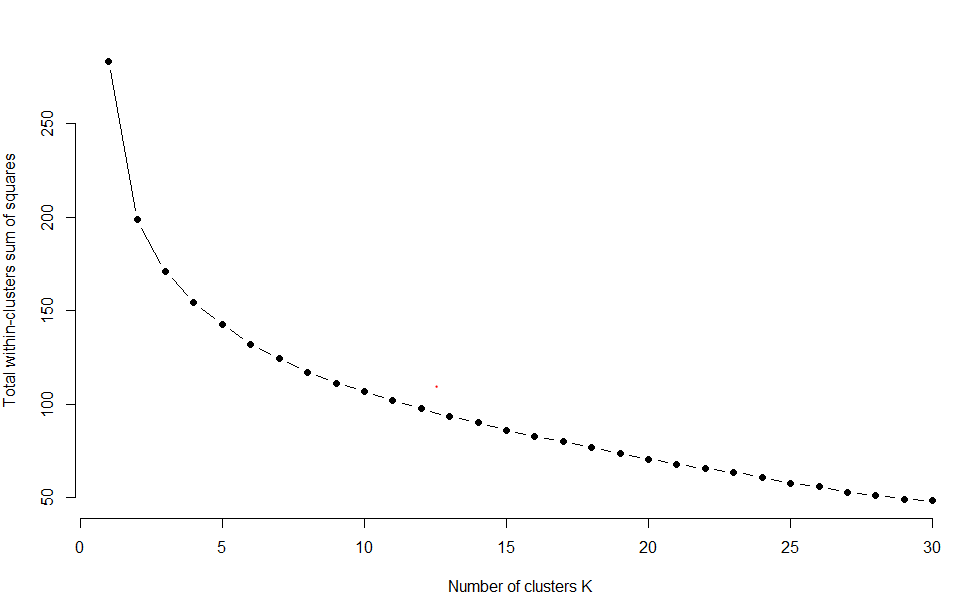

*Supplementary Figure S3*. Sum of squared error reduction with increased number of K clusters in K-means clustering. This plot is used for the “elbow-method” to determine number of clusters of an expression matrix.

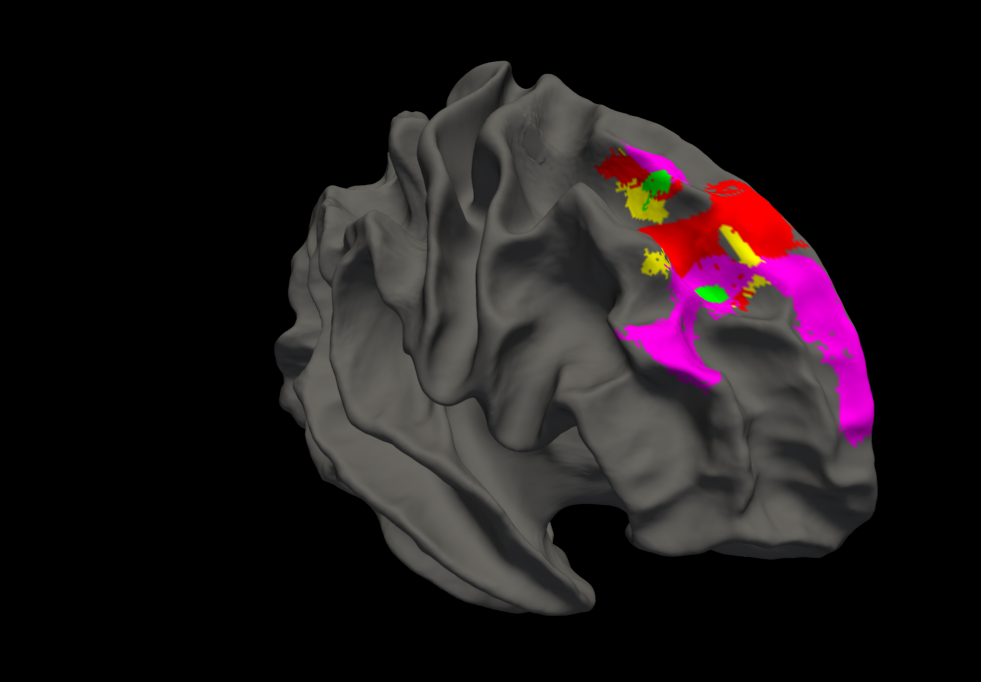

*Supplementary Figure S4.* Overlap across the split half replication in the right hemisphere. for the Inventory of Callous and Unemotional traits (ICU) and Center for Epidemiological Studies Depression scale (CESD). Yellow = CESD sample A, Green = CESD sample B, red = ICU Sample A, purple = ICU sample B. The more frontal clusters were all positively associated contiguous clusters and the more posterior clusters were all negative. These clusters overlap with the results from the full sample and are used to establish our “high confidence” clusters. The only split half-replicated results were in the right hemisphere.

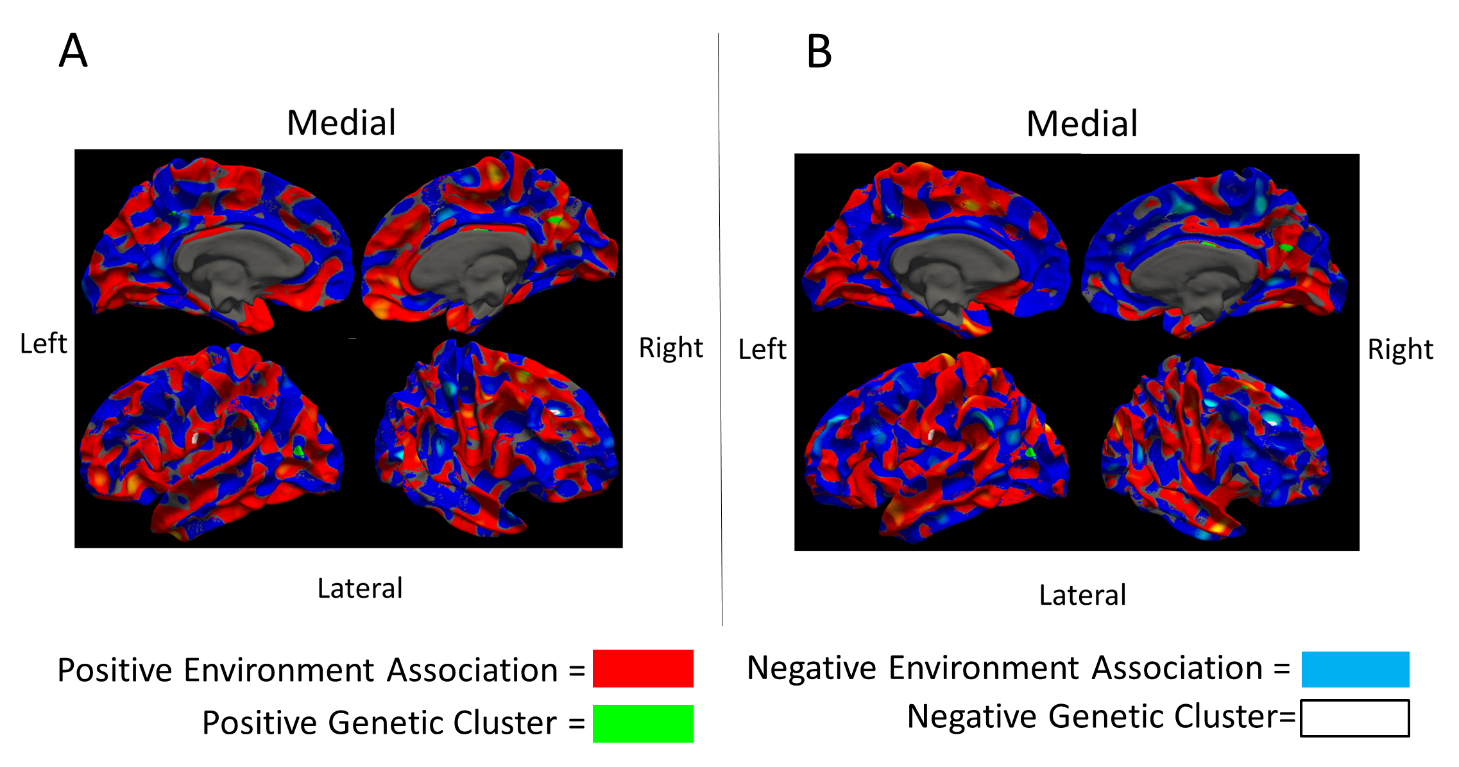

*Supplementary Figure S5.* Environmental heatmap of Center for Epidemiological Studies Depression scale (CESD; Panel A) and Inventory of Callous and Unemotional traits (ICU: Panel B), with genetic overlap clusters visualized over the environmental association pattern. Positive environmental associations are red; negative are blue. The genetic clusters from our main analysis show the positively associate clusters in white and the negatively associated clusters in green. None of the genetic and environmental associations were in the same direction on this map.

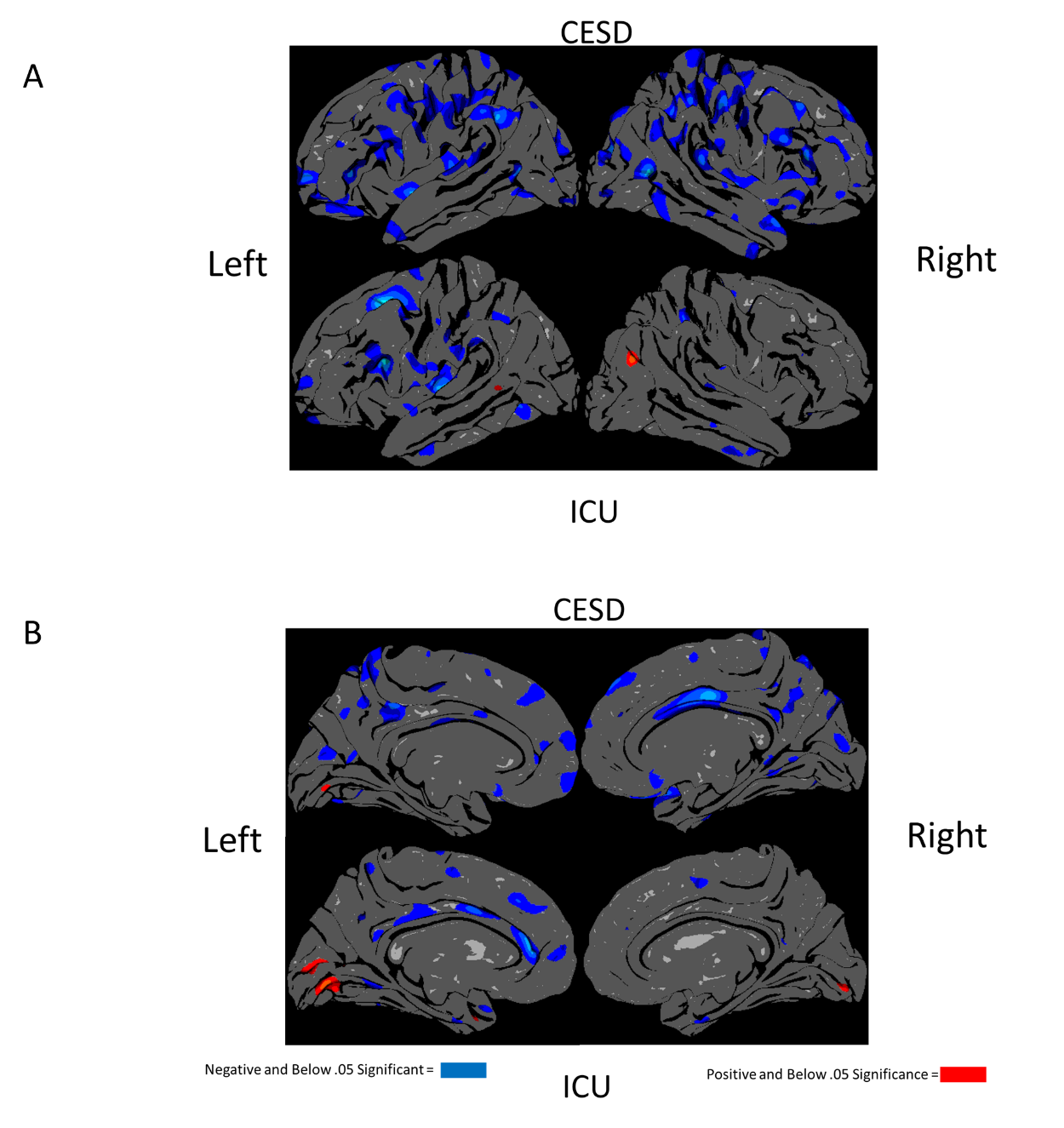

*Supplemental Figure S6.* Phenotypic brain map results a standard qdec Freesurfer analysis of the Center for Epidemiological Studies Depression scale (CESD) and Inventory of Callous and Unemotional traits (ICU). Lateral views of the brain are in Panel A and medial views are in Panel B. Clusters represent significance at nominal significance (*p* < .05). We chose a liberal threshold for this plot so we would have more likelihood of finding similarity between phenotypic and genetic clusters, as a key premise of our method is that they might differ.
